# Is *Anopheles gambiae* a natural host of *Wolbachia*?

**DOI:** 10.1101/491449

**Authors:** Ewa Chrostek, Michael Gerth

**Affiliations:** Max Planck Institute for Infection Biology, Berlin, Germany; Institute of Integrative Biology, University of Liverpool, United Kingdom

## Abstract

*Wolbachia* (*Alphaproteobacteria, Rickettsiales*) is an intraovarially-transmitted symbiont of insects able to exert striking phenotypes, including reproductive manipulations and pathogen blocking. These phenotypes make *Wolbachia* a promising tool to combat mosquito-borne diseases. Although *Wolbachia* is present in the majority of terrestrial arthropods, including many disease vectors, it was considered absent from *Anopheles gambiae* mosquitos, the main vectors of malaria in sub-Saharan Africa. In 2014, *Wolbachia* sequences were detected in *A. gambiae* samples collected in Burkina Faso. Subsequently, similar evidence came from collections all over Africa, revealing a high *Wolbachia* 16S sequence diversity, low abundance, and a lack of congruence between host and symbiont phylogenies. Here, we reanalyze and discuss recent evidence on the presence of *Wolbachia* sequences in *A. gambiae.* We find that although detected at increasing frequencies, the unusual properties of these *Wolbachia* sequences render them insufficient to diagnose natural infections in *A. gambiae*. Future studies should focus on uncovering the origin of *Wolbachia* sequence variants in *Anopheles* and seeking sequence-independent evidence for this new symbiosis. Understanding the ecology of *Anopheles* mosquitos and their interactions with *Wolbachia* will be key in designing successful, integrative approaches to limit malaria spread. Although the prospect of using *Wolbachia* to fight malaria is intriguing, the newly discovered strains do not bring it closer to realization.

**Significance:** *Anopheles gambiae* mosquitos are the main vectors of malaria, threatening around half of the world’s population. The bacterial symbiont *Wolbachia* can interfere with disease transmission by other important insect vectors, but until recently it was thought to be absent from natural *A. gambiae* populations. Here, we critically analyze the genomic, metagenomic, PCR, imaging and phenotypic data presented in support of the presence of natural *Wolbachia* infections in *A. gambiae.* We find that they are insufficient to diagnose *Wolbachia* infections and argue for the need of obtaining robust data confirming basic *Wolbachia* characteristics in this system. Determining *Wolbachia* infection status of *Anopheles* is critical due to its potential to influence *Anopheles* population structure and *Plasmodium* transmission.

## Introduction

*Wolbachia* is an obligate intracellular, intraovarially-transmitted bacterium living in symbiosis with many invertebrates (1). Depending on host and symbiont genotypes, and environmental conditions, *Wolbachia* has been shown to either affect the biology of its hosts in striking ways or exert only mild phenotypes. Some of the conspicuous *Wolbachia* phenotypes include reproductive manipulations, where maternally inherited symbionts favor survival and reproduction of transmitting females over non-infected females and non-transmitting males (2). One of the reproductive manipulations, cytoplasmic incompatibility (CI) (3), has been proposed as a tool to suppress mosquito populations and decrease arbovirus burden on humans (4, 5). Bidirectional CI - the inability of females to produce offspring with males harboring a different *Wolbachia* strain - has been successful in eliminating the filariasis vector, *Culex pipiens fatigans* from *Okpo*, Myanmar in 1967 (5), and suppressing *Aedes albopictus*, vector of dengue, Zika and West Nile virus, in trials in Lexington, Kentucky, California, and New York, USA (https://mosquitomate.com).

*Wolbachia* can also provide infected individuals with fitness benefits: nutrient provisioning (6), increase in reproductive output (7), and protection against pathogens (8, 9). The latter is also being used to eliminate vector-borne diseases. *Aedes aegypti* mosquitos artificially transinfected with protective *Wolbachia* are being deployed as a strategy to eradicate dengue virus (10–15). The data from one of the first release sites in Australia suggest that this strategy may limit the number of dengue cases in humans (15).

Malaria is a mosquito-borne disease that threatens around half of the world’s population (16). The potential for the use of *Wolbachia* to block malaria has been recognized since the symbiont’s anti-viral and anti-parasitic properties were first demonstrated in other insects (8–10, 17). However, *Anopheles* mosquitos were for long considered inhospitable for *Wolbachia* (18–20). This has started to change in 2006, when *Wolbachia* infections in *Anopheles* cultured cells were established for the first time (21). Next, transient somatic infections were created by intrathoracic inoculation of virulent *w*MelPop *Wolbachia* into adult mosquitos (22). In somatic transinfections, *Wolbachia* does not infect the germline (23), which is necessary for its maternal transmission and pathogen blocking-based field applications. Therefore, a successful generation of stable *Wolbachia* infections in *Anopheles stephensi* by Bian *et al.* was a big step towards field applications (24). Subsequently, gut microbiota of *A. stephensi and A. gambiae* was shown to hinder establishment of heritable *Wolbachia* infections in these species, and curing *Anopheles* of its microbiota enabled *Wolbachia* persistence (25). In 2014, the first evidence for natural *Wolbachia* infections was found in *Anopheles gambiae* and *Anopheles coluzzii* (two sibling mosquitos species of *Anopheles gambiae* species complex, considered the main malaria vectors in Sub-Saharan Africa – see Supplementary File 1 for details) from Burkina Faso (26). This was striking, as the natural *Wolbachia* phenotypes could change mosquito biology, population structure and, as such, affect malaria transmission. Several similar reports identifying *Wolbachia* sequences in *A. gambiae* populations across Africa have shortly followed (27–31).

Here, we examine the evidence on natural *Wolbachia* infections in *Anopheles gambiae* mosquitos and screen 1000 *Anopheles* genomes (Ag1000G) project data (32) to reveal that *Wolbachia* reads are extremely rare in this rich and randomized dataset. We re-analyze the data from which a genome of the putative *Wolbachia* endosymbiont of *Anopheles gambiae* was assembled (33) to show that the majority of reads in the sample originate from known *Wolbachia* hosts different than *Anopheles gambiae*. Finally, we discuss the requirements to diagnose a *Wolbachia* infections in a species previously considered uninfected, the potential ecological interactions which could lead to the observed *Wolbachia* sequence prevalence patterns, and their relevance for the design of successful, integrative approaches to limit malaria spread.

## Molecular evidence for natural *Wolbachia* in *Anopheles gambiae*

The first evidence of natural *Wolbachia* infections in malaria vectors comes from a study on field collected samples of *Anopheles gambiae* from Burkina Faso (26), in which *Wolbachia* sequences were detected through 16S V4 amplicon sequencing and a *Wolbachia*-specific PCR targeting the 438 bp ‘wSpec’ region of the 16S rDNA sequence (34). Furthermore, whole genome shotgun sequencing of two ovarian samples was performed. Out of over 164.6 million high quality *Anopheles*-depleted sequences obtained from two Illumina HiSeq lanes, 571 reads mapped to *Wolbachia* genomes, corresponding to a *Wolbachia* genome coverage of ∼0.05x. Overall, out of an average of over 1000 *Wolbachia* genes, only 134 had at least one read assigned to them. Moreover, 76 of the 571 reads mapped to *Wolbachia* transposases (26). This demonstrates that the *Wolbachia* sequences in these samples were of extremely low titer - the ratio of *Wolbachia* to host coverage was ∼1:4700. For comparison, in various *Drosophila melanogaster* sequencing projects, observed ratios ranged from 27:1 to 1:5 (35). The data described above represent the only genomic evidence for the presence of *Wolbachia* in *A. gambiae*.

To identify additional *Wolbachia* sequences in *A. gambiae*, we screened data generated in the Ag1000G project, which investigates genetic variance and population biology of *A. gambiae* (https://www.malariagen.net). We used the data released in the course of ‘phase 1 AR3’, namely Illumina sequences of 765 wild caught mosquitos from eight African countries (32). Reads for all samples were downloaded from the European Nucleotide Archive (ENA), and mapped to *Wolbachia* reference genomes. Using the criteria of Baldini *et al.* 2014 (26) (see Supplementary File 1 for details), we identified 446 reads from 96 libraries as matching to *Wolbachia*. In total, there were ∼7.89×10^10^ reads across 765 libraries, so only 1 in ∼150 million reads maps to *Wolbachia* (Fig. 1). This corresponds to less than one *Wolbachia* read per sequencing library on average, and, based on a large and broad sampling, provides independent evidence for only very sporadic presence, extremely low titer, or even absence of *Wolbachia* in *A. gambiae*.

**Figure 1.**
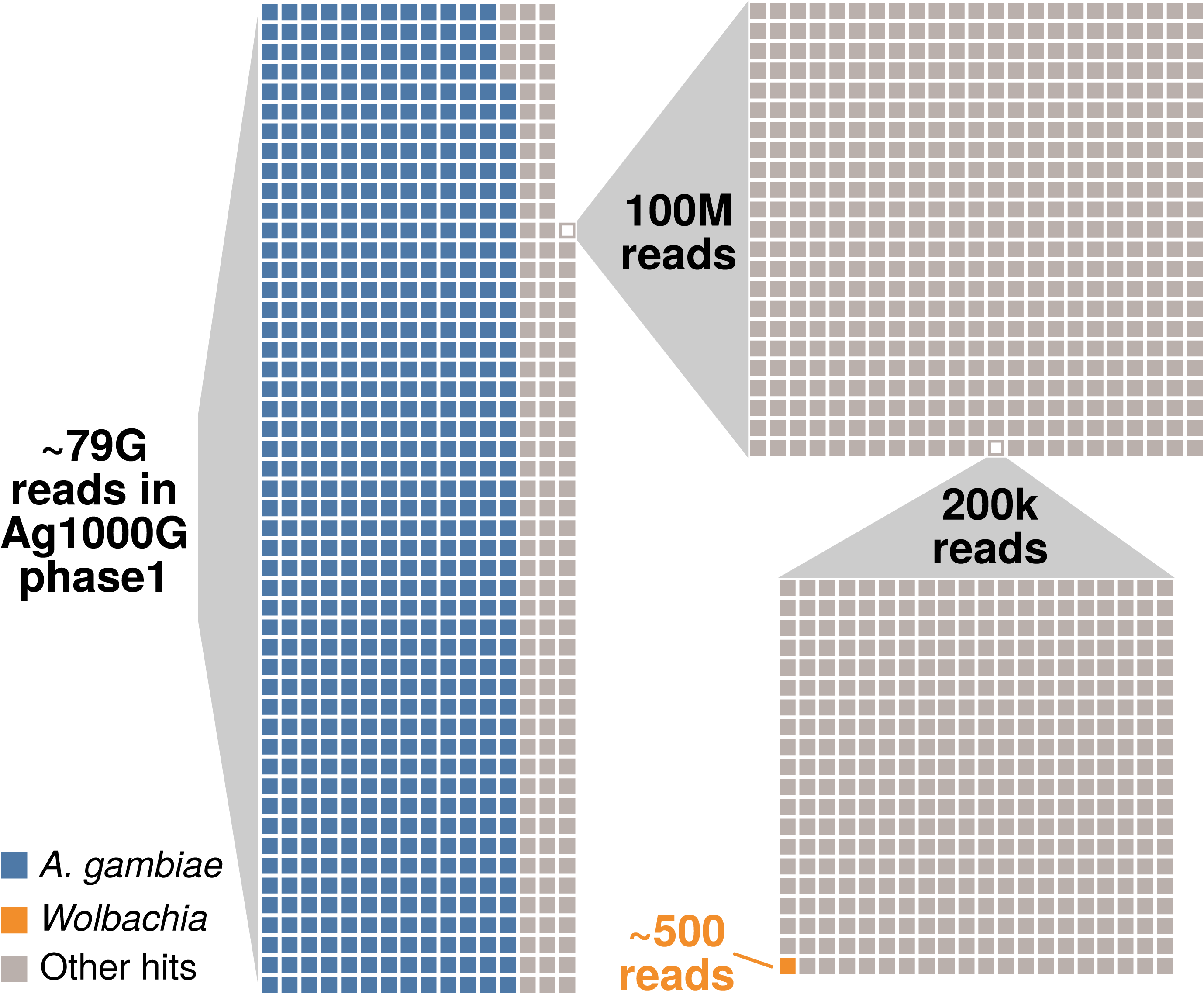
Taxonomic composition of the reads generated in the phase 1 of the Ag1000G project. In total, around 79 billion reads were generated from 765 *A. gambiae* mosquitos (32). Around 80% of these reads map to the *A. gambiae* host genome (represented by blue squares on the left). Panels on the right represent sequential magnifications of the portion of non-*Anopheles* reads, to visualize the proportion of reads mapping to *Wolbachia*.

Contrasting with our findings, a recent *in silico* screen of archived arthropod short read libraries extracted a highly covered *Wolbachia* supergroup B genome from a sample annotated as *A. gambiae* (33). To understand the reasons for this discrepancy we inspected the sequencing libraries used by Pascar and Chandler (33) and discovered that they contain a mix of sequences of several other potential *Wolbachia* hosts (Fig. 2). Based on the analysis of the ITS2 and COI haplotypes of the most abundant sequences (36, 37) we conclude that the assembled *Wolbachia* genome likely originates from *Anopheles* “species A” and not *A. gambiae* (Fig. 2, Fig. S1, Supplementary File 1). Our interpretation is in line with a recent discovery of a highly prevalent supergroup B *Wolbachia* strain, distinct from other supergroup B strains, in *Anopheles* “species A” (31). Our phylogenomic reconstructions further support this, as they place the newly assembled *Wolbachia* genome (33) within supergroup B, but separate from most other strains of this lineage (Fig. S1C). These analyses show that unambiguous identification of *Anopheles* species is an additional difficulty in detecting *Wolbachia* infections based on the sequencing data. Therefore, the newly reported genome does not contribute to the understanding of the elusive low titer *Wolbachia* naturally associated with *A. gambiae*.

**Figure 2.**
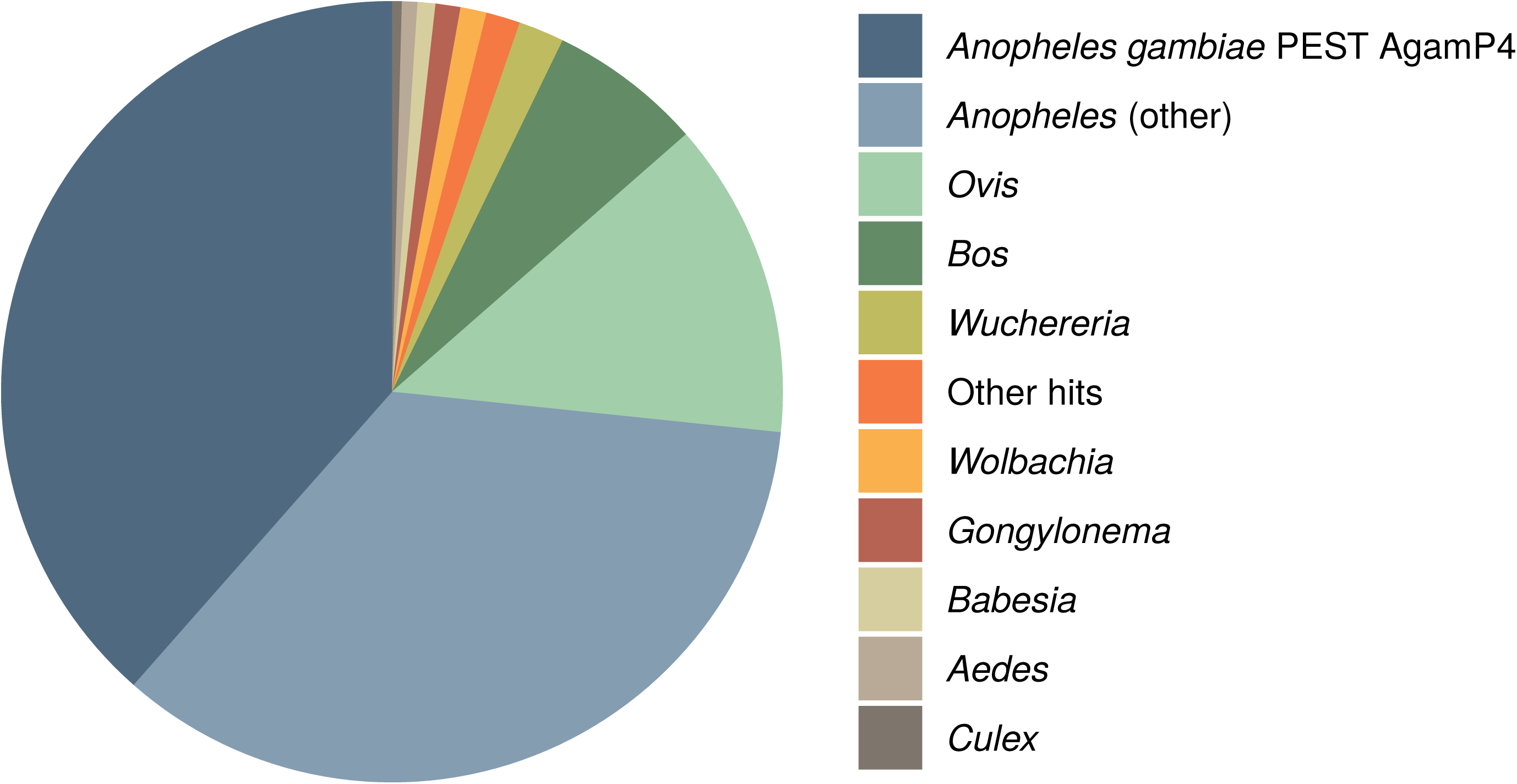
Taxonomic classification of reads in the libraries from which the genome of a putative *Wolbachia* symbiont of *A. gambiae* was assembled (BioSample SAMEA3911293). For more details, refer to Supplementary File 1 and Fig. S1.

The putative low titer *Wolbachia* infections required improved diagnostics. This has prompted Shaw *et al.* to modify the wSpec PCR protocol by including a nested pair of primers and increasing the number of cycles to 72, potentially amplifying the initial 16S template over 10^21^ times (28). The protocol was used in several subsequent studies (29–31), but proved unreliable, as Gomes *et al.* reported 19% of the technical replicates yielding discordant results, even when total number of cycles was increased to 80 (29). At the same time, the wSpec amplification protocol was sensitive enough to detect *Wolbachia* in a filarial nematode residing within one of the *Anopheles coustani* guts (30). Thus, this diagnostic test can detect *Wolbachia* in organisms interacting with *Anophele*s.

Meanwhile, Gomes *et al.* based their work on a 40-cycle qPCR-based assay (29). The robustness of this test is not clear, as no raw data were included. Other methods routinely used to detect low titer *Wolbachia* in insects, like PCR-southern blot or amplification of repeated sequences (e.g. the transposases with the highest coverage in genomic data) were never tested on *Wolbachia* sequences found in *Anopheles* (38, 39). Amplification of other *Wolbachia* sequences from putatively infected mosquitos, including *Wolbachia* surface protein and MLST genes, has also been challenging (26, 27, 29–31), requiring protocol modifications (30) or the use of more than one mosquito sample (31), and was unsuccessful in some cases (26, 27). Overall, detection of *Wolbachia* sequences in *A. gambiae* by PCR-based methods remains challenging.

In summary, very little sequence data is available for the putative *Wolbachia* symbiont of *A. gambiae*, despite several attempts of generating and extracting such data. One common feature of all of them is an extremely low titer, at the limit of detection of PCR-based methods. Even from the little data available, it is obvious that there is no single *Wolbachia* strain associated with *Anopheles gambiae* (Fig. 3). In fact, almost every *Wolbachia* 16S amplicon and sequence attributed to *A. gambiae* is unique, and their diversity spans at least two *Wolbachia* supergroups (genetic lineages roughly equivalent to species in other bacterial genera, Fig. 3) (40). In combination, we interpret the very low titers and the conflicting phylogenetic affiliations of the sequenced strains as incompatible with the notion of a stable, intraovarially-transmitted *Wolbachia* symbiont in *A. gambiae*. However, this conclusion requires alternative explanations for the presence of *Wolbachia* DNA in these malaria mosquitos.

**Figure 3.**
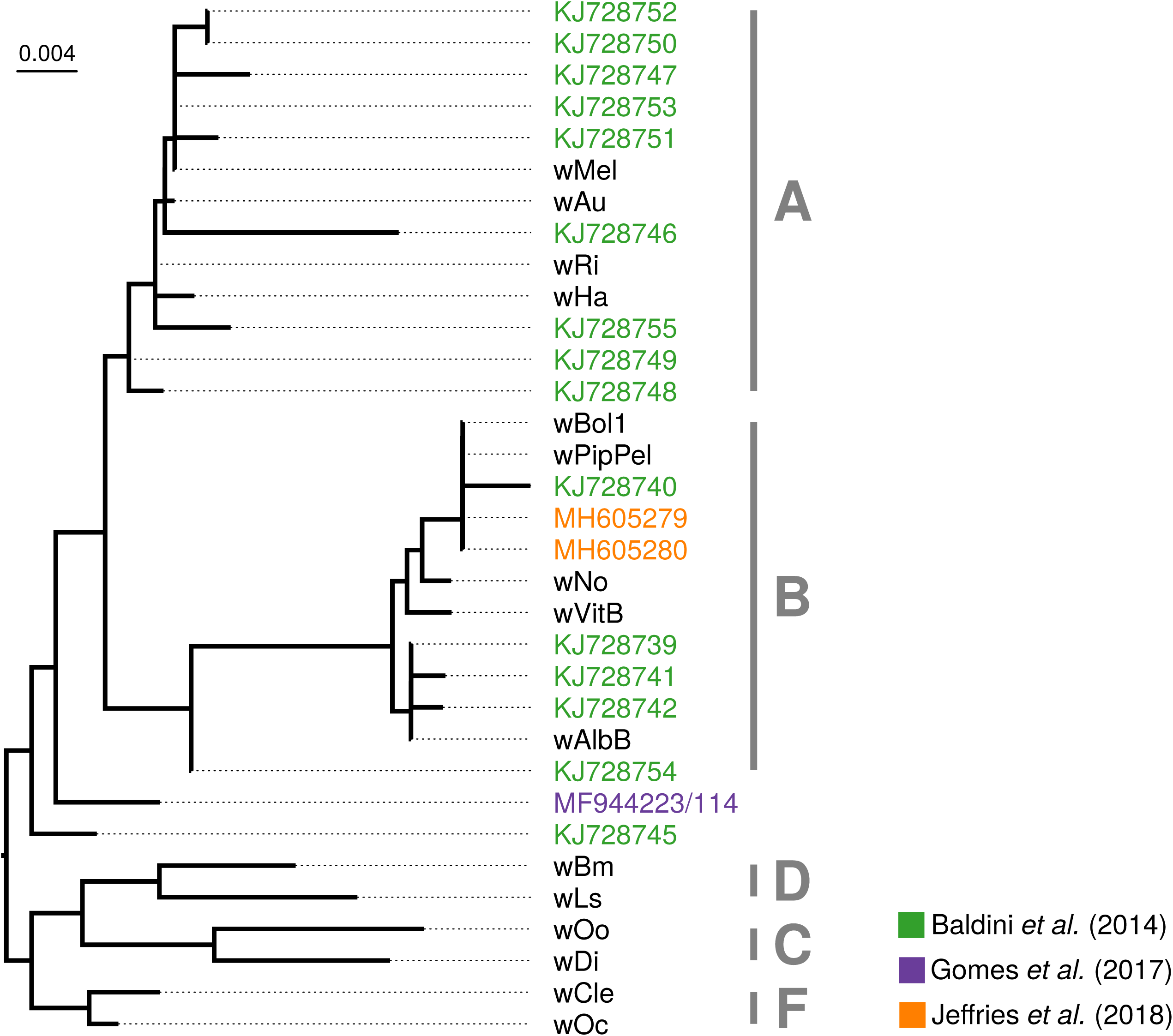
Phylogenetic placement of *Wolbachia* sequences from *Anopheles gambiae* based on 16S rRNA sequences. Alignment was done with Mafft using the ‘--auto’ option. Maximum likelihood tree was inferred with automatic model selection in IQ-TREE version 1.62 (60). Origin of sequences is indicated by colors (see legend), and tip names correspond to NCBI accession numbers. All other sequences are reference *Wolbachia* strains. Tentative supergroup affiliations are denoted with capital letters. Please note that the two *Wolbachia* 16S sequences determined by Gomes *et al.* are overlapping. Because the 117 bp overlap region is 100% identical between these two sequences we have merged them prior to phylogenetic analysis.

## Origin of *Wolbachia* sequences in *Anopheles gambiae*

The presence of *Wolbachia* DNA in *A. gambiae* samples could be explained not only by a stable *Wolbachia-Anopheles* symbiosis but also in several alternative ways. First, the signal could stem from *Wolbachia* DNA insertion into an insect chromosome (26). Fragments of *Wolbachia* genomes are frequently found within insect genomes (41–43), and the most spectacular cases include a nearly complete genome insertion in *Drosophila ananassae* (44). This possibility was discussed by Baldini *et al.*, but as the authors point out, the presence of the sequences only in some tissues, and the very low titer argue against this hypothesis (26). The second possibility discussed by Baldini and colleagues is the insertion of *Wolbachia* fragment into the chromosome of another, so far unidentified, mosquito-associated microorganism. However, this hypothesis does not help to explain the diversity of *Wolbachia* 16S sequences found in *Anopheles.*

Another hypothesis explaining the presence of *Wolbachia* sequences in *Anopheles* tissues would be contamination of the mosquito surface or gut. This contamination could come from several sources. First, ectoparasitic mites or midges, and endoparasitic nematodes in *Anopheles* could contaminate whole tissue DNA extracts, as shown by the detection of the *Wolbachia* symbiont of *Dirofilaria immitis* in *Anopheles coustani* DNA preparation (30). However, the presence of unknown symbionts or parasites with novel *Wolbachia* strains is very challenging to test for.

The second possible source of *Wolbachia* contamination are plants. It has been shown that *Wolbachia* can persist in plants on which *Wolbachia*-infected insects feed, and then be detected in previously uninfected insects reared on the same plant (reviewed in (45)). As malaria vectors feed on plant nectar and fruits in the wild, *Wolbachia* DNA traces from these sources could accumulate in their guts. Feeding on *Wolbachia* infected food could explain *Wolbachia* 16S rDNA encounter in the ovaries, as adjacent gut can easily be perforated during dissections, releasing content and contaminating other tissues. Again, *Wolbachia* sequences from the gut could also explain detection of *Wolbachia* sequences in larvae, as eggs and larval habitats could be contaminated with adult feces.

Another possible source of contamination are other insects co-habiting the collection sites. *Culex, Aedes* and *Anopheles* species can be found in sub-Saharan Africa, and all genera include natural *Wolbachia* hosts. This route of contamination seems especially plausible for mosquito larvae, which are avid predators, attacking other water inhabiting insects. Moreover, *Wolbachia* 16S sequences can be detected in the water storage containers inhabited by larvae of various mosquito species (Supplementary File 1), and as such could also be acquired by newly emerging adults and females during egg laying (46). Unfortunately, we have no data on the water composition of the breeding sites of the putative *A. gambiae Wolbachia* carriers, which could explain *Wolbachia* sequences presence across the mosquito life cycle.

The data on natural *Wolbachia* infections in *A. gambiae*, together with similar reports suggesting *Wolbachia* infections in species previously considered uninfected, e.g. *A. stephensi* (47), *A. funestus* (48) and *A. aegypti* ((47, 49, 50) but also (51, 52)), should be carefully examined, as all have aquatic, detritus-feeding and predatory larvae, while adults are terrestrial and can feed on nectar. Thus, bacteria and/or contaminating sequences could spread between these and other organisms sharing the same niches, necessitating studies designed to discern candidates for symbiotic taxa from transient and contaminating bacteria. Sampling of the mosquitos along with their environments and co-habiting species may help to reveal the origin and nature of *Wolbachia* sequences identified in *A. gambiae*.

Importantly, the contamination from any of the mentioned sources cannot be ruled out with the data currently available. The previously mentioned sequencing of two *Wolbachia*-positive ovary samples resulted in 571 (out of ∼800,000,000) reads classified as *Wolbachia* (0.000063%) (26). For a highly sensitive sequencing technique such as Illumina sequencing, this falls well within the expected coverage of contaminants. Deep shotgun sequencing of eukaryotes usually results in some non-target sequences from environmental contaminants, and it is unlikely that the *A. gambiae* libraries are an exception (53–55). Contamination stemming from non-target microbial taxa is especially problematic in low biomass samples (56), such as single mosquito ovaries. Adding to the difficulty, all of the studies reporting *Wolbachia* from amplicon or metagenomic sequencing do not present negative controls (e.g. sequencing of extraction or blank controls, quantification of microbial taxa, sequencing of mock communities (26, 27, 29–31)). This is not to say that the *Wolbachia* sequences definitely constitute contaminants, but they are simply not discernible from such. In general, the detection of very low titer *Wolbachia* through highly sensitive methods (nested PCRs, Illumina sequencing) alone is not sufficient to conclude that an intracellular, inherited symbiont is present in a sample.

## Expected features of natural *Wolbachia* from *Anopheles gambiae*

While sequence data alone are insufficient to determine if *Wolbachia* is a symbiont of *Anopheles gambiae*, and assembly of complete genomes has not been achieved due to low sequence abundance, other hallmarks of symbiotic interactions between the taxa can be used to support this claim.

First, intracellular localization is imperative for *Wolbachia*. The only published image of natural *Wolbachia* infections from *A. gambiae* is an indirect fluorescence *in situ* hybridization, using Cy3-labelled probe, anti-Cy3 mouse antibody, and anti-mouse Alexa448 secondary antibody (see Fig. 1 in Ref (28)). The probe was designed to hybridize within, by then, the only PCR-detectable *Wolbachia* sequence - the wSpec amplicon region. However, the low resolution of the image and the lack of host membrane staining do not allow to confirm the wSpec intracellular localization (28). Electron microscopy showing an immunogold-labelled *Wolbachia*, or a high-resolution FISH combined with a membrane staining would provide unequivocal visual evidence for the existence of intracellular *Wolbachia* infections in *A.* g*ambiae*.

Second, *Wolbachia*’s intracellular lifestyle is directly related to its mode of transmission, which is expected to occur from mother to offspring within the mother’s ovaries. In the first study on natural *Wolbachia* in *A. gambiae*, maternal transmission of the detected wSpec sequences was also examined. In this experiment, five wSpec-positive wild-collected gravid females oviposited in the lab and their larval progeny was tested for wSpec amplification (detected in 56% to 100% of the offspring) (26). However, intraovarial transmission of *Wolbachia* was never explicitly addressed. Surface sterilization of eggs after oviposition would help to determine the transmission mode of these sequences, just as testing for and excluding horizontal (between larvae or adult to larvae) and paternal wSpec sequence transmission. These experiments would help to confirm that *A. gambiae* is infected with an intracellular, transovarially transmitted symbiont and, together with the PCR evidence, diagnose a stable *Wolbachia* infection.

## *Wolbachia* symbionts of *Anopheles gambiae* and malaria

*Wolbachia* phenotypes similar to those observed in other insect hosts could have a huge impact on wild *Anopheles* populations and malaria transmission. Reproductive manipulations and fitness benefits could increase the proportion of biting females spreading the disease, while pathogen blocking could limit *Plasmodium* prevalence in the wild mosquito populations. Understanding *Anopheles gambiae* biology is crucial for the design of effective strategies aiming at limiting *Plasmodium* transmission.

Targeted *Wolbachia-*based *Plasmodium* control strategies, similar to the ones used for dengue and Zika virus control, are another exciting prospect. However, they are not reliant on *Wolbachia* symbionts naturally associated with *Anopheles.* Insect populations could equally well be suppressed by the release of males carrying incompatible *Wolbachia* strains by bidirectional CI on infected population or by unidirectional CI on an uninfected one. The same applies to *Wolbachia*-induced pathogen blocking. Existing initiatives to control dengue and Zika virus with *Wolbachia*-conferred antiviral protection use naturally uninfected *Aedes aegypti* mosquitos that were artificially transinfected with *Wolbachia* from a different insect species (12). These mosquitos benefit not only from protection by the core and yet unknown mechanism, but also from immune system upregulation caused by a recent transinfection with *Wolbachia* (10). Thus, the *Wolbachia*-based population suppression and disease blocking can work in species not commonly infected with *Wolbachia* in the wild.

The presence and, subsequently, *Plasmodium* blocking properties of the presumed natural *Wolbachia* strains in *A. gambiae* remain to be confirmed. Given that *Wolbachia* detection in *A. gambiae* remains challenging (with PCR-based replicate experiments yielding discordant results (29)), it was surprising that two studies have reported negative correlations between the low titer *Wolbachia* sequences and *Plasmodium* (28, 29). As pathogen protection has been shown to depend on the symbiont titer (57–59) and has so far only been detected in strains exhibiting relatively high bacterial load, it is likely to be absent in *A. gambiae* (31). Moreover, CI necessary for the spread of *Wolbachia* in artificially infected vector populations was also not detected (28). Reliable protocols for the detection of *Wolbachia* in *A. gambiae*, together with independent repetition efforts seem necessary to characterize the potential of the putative *A. gambiae* symbionts for their deployment in vector or disease control programs.

In summary, although using *Wolbachia* to fight malaria has been eagerly anticipated, naturally occurring *Wolbachia* strains in *Anopheles* were never an absolute requirement for this to be successful. Even now, their presence, phenotypes and suitability for deployment in disease control remain to be confirmed. However, they should be studied, as understanding *Anopheles gambiae* biology and ecology, including its interactions with other micro- and macroscopic organisms, is crucial for designing effective malaria elimination programs.

## Conclusions

The evidence for natural *Wolbachia* infections in *Anopheles gambiae* is currently limited to a small number of highly diverse, very low titer DNA sequences detected in this important malaria vector. Further efforts towards characterization of the interaction between *Wolbachia* sequences and *A. gambiae* are required to establish that this is a true symbiotic association. Demonstrating the presence of intracellular bacterial cells and their intraovarian transmission are prerequisites to diagnose a symbiosis. Additionally, genomic data could shed light on the features of these *Wolbachia* and may reveal the origin of the sequences and the ecological interactions that caused their acquisition by *A. gambiae* mosquitos. Finally, ascertaining phenotypes associated with these *Wolbachia* sequence variants will improve our understanding of *Anopheles gambiae* biology, and as such inform future strategies aimed at limiting malaria spread and eventual disease eradication.

The fact that *Wolbachia* sequences were encountered multiple times by independent groups of researchers clearly indicates present or past, direct or indirect ecological interaction between *Wolbachia* and *Anopheles gambiae* across Africa. While in-depth investigations of these interactions will be interesting from a basic biology, evolutionary, ecological and disease control perspective, current data indicate that the postulated natural *Wolbachia* infections in *Anopheles* will be of limited use for application in fighting malaria with *Wolbachia*.

## Supporting information

## Acknowledgements

We thank Francis Jiggins for a suggestion on using the *Anopheles gambiae* 1000 genomes project data and for the critical reading and helpful comments on the manuscript. We also thank Elena Levashina and Elves Heleno Duarte for comments on the draft of this work.

EC was supported by EMBO Long-Term Fellowship (EMBO ALTF 1497-2015), co-funded by Marie Curie Actions by the European Commission (LTFCOFUND2013, GA-2013-609409) and FEBS Long Term Fellowship. MG was supported by a Marie Curie Fellowship of the European Commission (H2020-MSCA-IF-2015, 703379).

## Supplementary figure legend

**Figure S1 |** Phylogenetic assessment of SAMEA3911293 taxonomic composition. A) Phylogenetic reconstruction of *Anopheles* species based on ITS2 alignment of previously published data and all ITS2 contigs present in the meta-assembly of all libraries from SAMEA3911293. Sequences recovered from this library are highlighted in blue. B) Phylogenetic reconstruction of *Anopheles* based on mitochondrial COI. Sequences from the SAMEA3911293 meta-assembly are highlighted in blue. C) Phylogeny of *Wolbachia* supergroup B based on concatenated core genome alignments of all strains with a (draft) genome in NCBI. Again, the strain isolated from SAMEA3911293 is highlighted in blue.

