## Supplementary figures and images for "Is *Anopheles gambiae* a natural host of *Wolbachia*?"

### Supplementary file 1

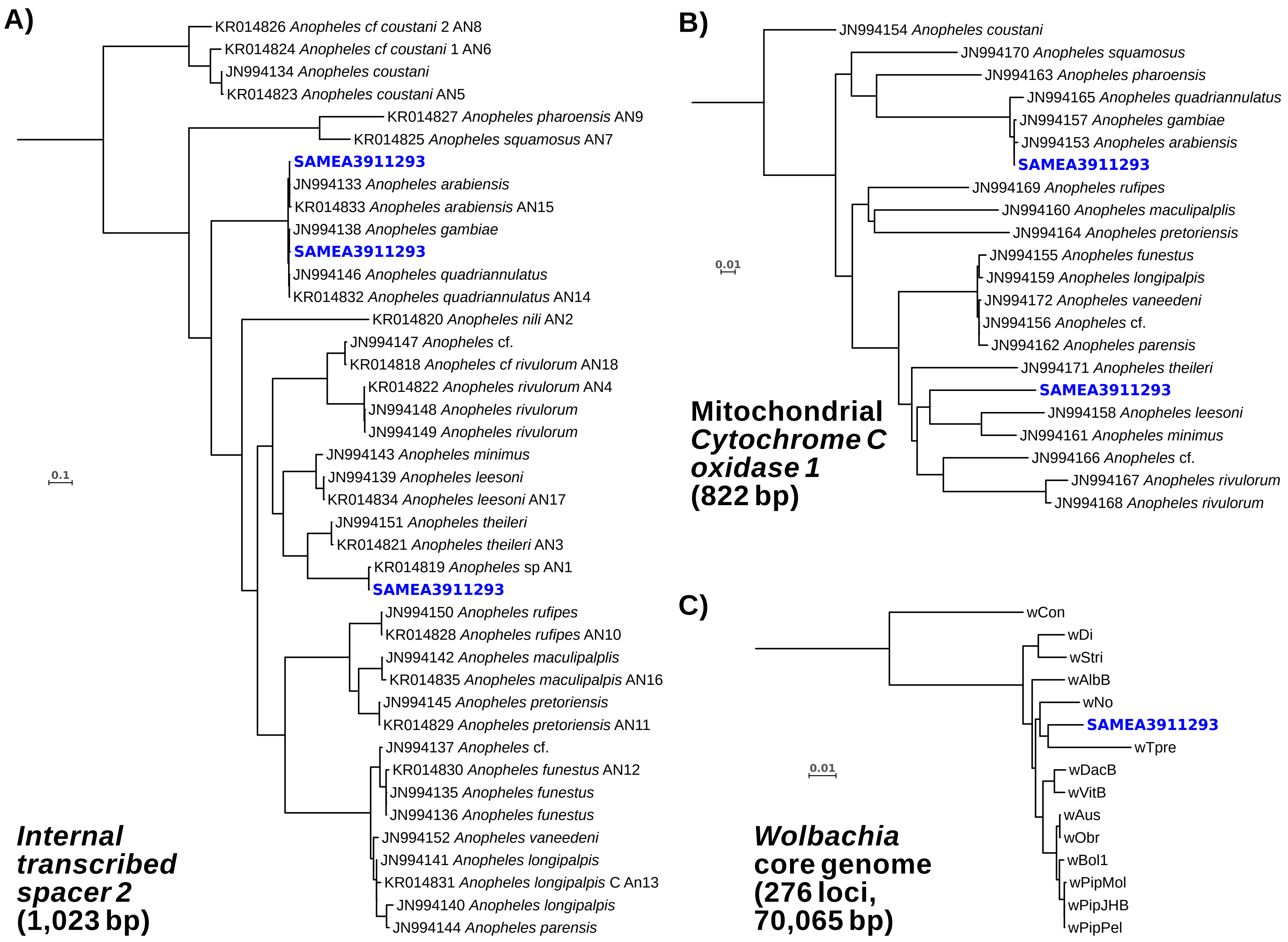
