## Supplementary material for "Is *Anopheles gambiae* a natural host of *Wolbachia*?"

**Supplementary file 1**

*Note on Anopheles gambiae taxonomy*

*Anopheles gambiae* is currently considered a species complex containing multiple distinct genetic lineages (White et al. 2011). Here we consider only *A. gambiae sensu stricto* (previously *A. gambiae* form S) and *A. coluzzi* (*A. gambiae* form M), as these are the only members of the species complex in which *Wolbachia* was detected and characterised. *Wolbachia* specific wSpec primers were also used to amplify fragments from *A. arabiensis* (which is also part of the *A. gambiae* complex, Shaw et al. 2016 and Jeffries et al. 2018). However, as no sequences are available to characterize these infections, here we have focused on the *A. gambiae* and *A. colluzzi* and we refer to these two species collectively as '*A. gambiae*'.

*Screening for Wolbachia in Ag1000G data*

To determine if *Wolbachia* sequences are commonly found in *Anopheles gambiae*, we screened data generated in the '*Anopheles gambiae* 1000 genomes' (Ag1000G) project. We downloaded the data of the phase 1 public release, which included Illumina sequences from 765 wild caught *A. gambiae* from the European Nucleotide Archive (https://www.ebi.ac.uk/ena/data/view/PRJEB18691). For each of the 765 samples, a single bam file was downloaded, and all fastq reads extracted using SAMtools version 1.9 (Li et al. 2009). The reads were then mapped to six complete *Wolbachia* genomes representing the major phylogenetic lineages of this genus (Table 1) using NextGenMap version 0.5.5 (Sedlazeck et al. 2013).

Table 1 List of *Wolbachia* genomes used in the screen

| **Strain** | **Native host** | **Supergroup** | **NCBI BioProject** | **Reference** |
|---|---|---|---|---|
| wMel | *Drosophila melanogaster* | A | PRJNA272 | Wu et al. 2004 |
| wPipPel | *Culex pipiens* | B | PRJNA30313 | Klasson et al. 2008 |
| wOo | *Onchocerca ochengi* | C | PRJEA81837 | Darby et al. 2012 |
| wBm | *Brugia malayi* | D | PRJNA12475 | Foster et al. 2005 |
| wFol | *Folsomia candida* | E | PRJNA299291 | Faddeeva-Vakhrusheva et al. 2017 |
| wCle | *Cimex lectularius* | F | PRJDB748 | Nikoh et al. 2014 |

In accordance with the genetic divergence expected within the genus (Chung et al. 2018), and to reduce spurious alignments, we discarded all reads with an identity lower than 95% to any of the references and also excluded alignments <50bp (<50% of the average read length). Next, we followed the protocol outlined by Baldini *et al.* (2014) to extract the reads that matched to *Wolbachia*:

1) All matches to ribosomal RNA genes were excluded.

2) All remaining reads were blasted against the NCBI 'nt' database using a word size of 7 and further filtered:

- We kept reads if they matched to any *Wolbachia* sequence with length >95bp and identity >80%, but only if there were no matches to other taxa with length >80bp;
- We also kept hits to *Wolbachia* with identity >90% and no match to other taxa with identities >80%.

*Analysis of NCBI BioSample SAMEA3911293*

In a recent *in silico Wolbachia* screen of many different short reads libraries from NCBI's SRA database, Pascar & Chandler (2018) detected a *Wolbachia* strain in one library (ERR1554906) annotated as *Anopheles gambiae* (NCBI BioSample accession SAMEA3911293). They have further isolated a fairly complete draft genome of this *Wolbachia* strain (Pascar & Chandler 2018). The computational pipeline employed by Pascar & Chandler (2018) was oriented towards automated detection and isolation of *Wolbachia* reads from short read libraries. While this may be a powerful approach to detect so far unrecognised *Wolbachia*-host associations, the pipeline did not include a number of quality and sanity checks. Importantly, Pascar & Chandler (2018) did not check the taxonomic classification of the libraries, i.e, if the library which contained *Wolbachia* actually stems from *A. gambiae*.

To confirm that this *Wolbachia* strain was isolated from *A. gambiae*, we downloaded all reads associated with the sample (three runs in total: ERR1554906, ERR1554870, and ERR1554834). It should be noted that this sample is also part of the Ag1000G project (see above), but was not publicly released yet. Furthermore, no metadata are available for this sample on NCBI (e.g., geographical origin, tissue used for DNA extraction, number of individuals pooled, etc). We mapped the downloaded reads to the *Anopheles gambiae* reference genome (strain PEST AgamP4 that is also used as a reference in the Ag1000G project) with NexGenMap as described above, but using the less sensitive default mapping options. Because the majority of reads did not map to this reference, we classified the remaining reads by:

1) Performing an assembly of all unmapped reads with MEGAHIT version 1.1.1-2-g02102e1 (Li et al. 2015);

2) Taxonomic classification of contigs of the resulting meta-assembly through BLAST+ (Camacho et al. 2009) searches against a local copy of the NCBI 'nt' database (e-value cutoff 1e-12, alignment length ≥100 bp, best match was used for taxonomic assignment);
3) Mapping of all reads not matching the *Anopheles gambiae* reference genome to the meta-assembly, and assigning the reads with the classifications of the contigs they mapped to.

The results of this classification are depicted in Figure 2 and Supplementary Figure S1. The majority of reads not mapping to the reference could be classified as different *Anopheles* species. Among other common taxa encountered in the sample are several potential *Wolbachia* hosts (*Culex*, *Aedes*, *Wuchereria*). This demonstrates that the investigated libraries were not constructed from a "pure" *Anopheles gambiae* sample, but rather from a pool of different host species (including *A. gambiae* and at least one other *Anopheles* species), potentially a metagenomic sample. Without metadata it is however not possible to determine how this sample was collected.

To identify other potential *Anopheles* species in the libraries, we performed phylogenetic analyses based on two markers commonly used in *Anopheles* species assignment: mitochondrial cytochrome C oxidase subunit 1 (COI) and internal transcribed spacer 2 (ITS2). We identified these fragments in the meta-assembly by BLAST searches using the corresponding *A. gambiae* sequences as query. After merging overlapping but otherwise identical matches, we found three and two distinct sequences for ITS2 and COI, respectively. Phylogenetic analyses were performed on alignments of these sequences together with reference sequences from previous phylogenetic studies on *Anopheles* (Lobo et al. 2015; Norris & Norris 2015). The corresponding phylogenetic reconstructions are depicted in Supplementary Figure S1A and B.

For both ITS2 and COI, we found haplotypes in the investigated libraries that clustered within the *A. gambiae* complex and therefore likely stem from *A. gambiae* or a very closely related species. However, for both loci we also recovered a sequence that is only very distantly related to *A. gambiae*. In the ITS2 tree, the sequences are almost identical to a sequence from a presumably undescribed *Anopheles* species, denominated "species A" in Stevenson et al. (2012) and "N1" in Lobo et al. (2015). In the COI tree, there is no very close match to the haplotype from the short read library, but it is evident that *A. gambiae* is only distantly related. Closer inspection of our metaassembly revealed the presence of a single contig spanning the complete mitochondrial genome of this species. Online BLAST searches against the NCBI database showed that it is only ~92% identical to the closest mitochondrial genome in the database (*A. stephensi*) and only 91% identical to the mitochondrial genome of *A. gambiae*.

These findings, together with the taxonomic classification of the reads in this sample discussed above, strongly suggest that in addition to *A. gambiae*, there is at least one other *Anopheles* species (most likely "species A") present in the sample SAMEA3911293. Because *Wolbachia* was not detected in our screen of 765 *A. gambiae* samples, we think that it is very likely that the *Wolbachia* sequences in this sample stem from the *Anopheles* "species A", or even a third species (e.g., *Culex* or *Aedes*) rather than from *A. gambiae*. Intriguingly, in the PCR based *Wolbachia* screen by Jeffries et al. (2018), *Anopheles* "species A" was found to be frequently infected with a *Wolbachia* strain of supergroup B that is distinct from other so far sequenced supergroup B strains. Our phylogenetic reconstruction of *Wolbachia* supergroup B based on core genome loci (Supplementary Figure S1C) further supports our interpretation, as the *Wolbachia* strain isolated from the investigated short read libraries clusters within *Wolbachia* supergroup B, but is distinct from other strains of this phylogenetic group.

*Screening for Wolbachia in amplicon data generated from water storage containers*

To assess the plausibility of *Wolbachia* sequences being obtained by *Anopheles* larvae from environmental sources, we explored a 16S amplicon dataset generated in a study investigating the bacterial composition of water storage containers with and without mosquito larvae (Nilsson et al. 2018). We downloaded the raw reads associated with this study from https://www.ncbi.nlm.nih.gov/bioproject/436283 and used NextGenMap to align all reads against a collection of *Wolbachia* 16S rRNA comprising all known supergroups of *Wolbachia,* which was taken from Glowska et al. (2015). We found hits in 9 out of 80 investigated libraries, including libraries constructed from water with and without inhabiting mosquitoes. Consensus sequences were created from all reads of a single library that matched *Wolbachia*, and each candidate *Wolbachia* sequence was blasted against the NCBI 'nt' database. All sequences matched a *Wolbachia* sequence in the database with at least 98% identity and 99% query coverage, confirming these sequences to be originating from *Wolbachia*.

This brief analysis demonstrates that *Wolbachia* sequences can be detected in amplicon sequences from environmental sources even if no apparent *Wolbachia* host is present. This highlights the importance of further verification of *Wolbachia* presence if determined with highly sensitive methods such as massively parallel amplicon sequencing, which are especially prone to contamination.
